# Rapidly evolving orphan immunity genes protect human gut bacteria from intoxication by the type VI secretion system

**DOI:** 10.1101/2025.05.03.651265

**Authors:** Amirahmad Azhieh, Paul Hernandez, Alexander C. Anderson, David Sychantha, Adrian J. Verster, John C. Whitney, Benjamin D. Ross

## Abstract

Bacteria encode diverse mechanisms for mediating interbacterial antagonism through the exchange of toxic effector proteins. Although the structure, function, and regulation of these pathways has been well established for many organisms, an understanding of their ecological and evolutionary dynamics lags behind. Type VI secretion systems (T6SS) deliver effectors between competing Gram-negative bacteria, including among mammalian gut Bacteroidales, resulting in the evolution of elaborate defense mechanisms that protect against T6SS attack. One such mechanism is the <u>r</u>ecombinase-associated <u>a</u>cquired interbacterial <u>d</u>efence (rAID) system, which harbors arrays of orphan immunity genes that diverge in sequence from T6SS-associated cognate immunity genes. It is not known if such sequence divergence impacts rAID orphan immunity function, or how rAID distribution across microbiomes relates to the T6SS. Here, we show that divergent rAID orphan immunity factors that possess SUKH domains allow bacteria to survive intoxication by cognate effectors. Such protection is due to high affinity protein-protein interactions between orphan immunity and effector that are comparable to that of cognate effector-immunity. Unlike other examples of T6SS effector-immunity interactions, we find that the binding interface is comprised of electrostatic interactions with a high degree of redundancy underlying its protective capacity. Finally, we quantify orphan immunity and effector gene abundance and dynamics across human gut metagenomes, revealing patterns of co-occurrence indicative of positive selection. Population genetic analyses of longitudinal data suggests that orphan immunity genes accumulate non-synonymous mutations that lie at the predicted effector-immunity interface. Together, our findings establish rAID orphan immunity genes as important bacterial fitness determinants in the human gut.

## INTRODUCTION

Understanding the ecological relationships between co-existing microbes and the respective evolutionary forces shaping their genomes is a fundamental goal of microbiology. This is particularly relevant for the mammalian gut where diverse populations of microbes engage in direct and indirect interactions with one another. However, the inaccessibility of the gut environment has led to limited insight into what molecular mechanisms underlie these interactions and there is speculation that our lack of understanding of these processes has hindered progress into the development of effective microbiota directed therapies [1]. Antagonistic interactions between co-resident bacteria are a common feature of many bacterial communities, including in the mammalian gut [2–4]. Among the pathways that mediate antagonism, the type VI secretion system (T6SS) is a contact-dependent protein translocation pathway encoded by many Gram-negative bacteria that exports bacteriolytic and bacteriostatic effector proteins [5]. When delivered to targeted cells, these effectors degrade conserved and essential molecules leading to increased fitness of producer cells in the context of bacterial competition during prolonged cell-cell contact [5]. Since many T6SSs indiscriminately target kin cells as well as non-kin cells, T6SS effector-encoding bacteria produce cognate immunity factors to prevent self-intoxication and these factors typically neutralize effector toxicity through occlusion of effector enzyme active sites via highly specific protein-protein interactions [6].

While many T6SSs are well-characterized at the molecular level, with notable advances in understanding its macromolecular structure and the mechanisms underlying diverse effector activities [7, 8], less is known about the role of these systems in the ecology and evolution of natural microbial communities. In the human gut microbiome, the dominant T6SS-encoding bacteria are derived from the highly prevalent and abundant order Bacteroidales [9–13]. These bacteria encode three distinct T6SSs, denoted <u>g</u>enetic <u>a</u>rchitecture (GA) 1-3 based on differences in gene content and syntenic organization, and each T6SS harbors extensive non-overlapping repertoires of effector and immunity genes [10–12, 14]. Of these T6SSs, GA3 systems are best understood likely owing to their exclusive existence in *Bacteroides fragilis*, a well-studied organism due to its ability to potently stimulate host immunity during development and, in rare cases, cause serious extraintestinal infections [15]. *B. fragilis* strains that utilize the GA3 T6SS exhibit a substantial fitness advantage over their competitors *in vitro* and in the context of co-colonization studies in mice [9, 13, 16, 17]. Comparatively less is known about the activity and function of the integrative and conjugative element (ICE)-encoded GA1 and GA2 subtypes. Unlike GA3, GA1 and GA2 exhibit extensive and varied patterns of occurrence in the genomes of different species of Bacteroidales [10, 11]. Within individual microbiomes, the horizontal spread of GA1 and GA2 systems between co-resident strains and species has been inferred by whole genome sequencing of bacterial isolates and metagenomic surveys [11, 12]. Bacteria acquiring one of these mobile T6SSs are expected to be protected from growth inhibition by T6SS activity due to the acquisition of cognate immunity genes, thereby providing an explanation for their wide dissemination [12, 14].

Bacteria that are the targets of T6SS-mediated antagonism are the subject of intense selective pressure and as a result have evolved defense mechanisms to withstand intoxication [4, 18, 19]. Some of these mechanisms appear to be general, such as elevation of stress response pathways or elaboration of exopolysaccharides. Others are highly specific and involve the accumulation of immunity genes predicted to neutralize the toxic activity of diverse effectors. Immunity genes that are harbored by a genome that lacks cognate effectors are referred to as orphan immunity genes and have been identified in diverse species including among gut Bacteroidales, which harbor several gene clusters involved in interbacterial defense that each encode distinct orphan immunity repertoires [16, 20]. For example, the acquired interbacterial defense-1 (AID-1) system is found in the genomes of some *Bacteroides* species including *B. fragilis* and *B. ovatus* and harbors GA3-specific immunity factors that are highly effective at neutralizing the toxicity of GA3 effectors *in vitro* and in mice [16]. In contrast to AID-1, the recombinase-associated AID (rAID) pathway is a widespread T6SS defense system that is broadly distributed across genera within the order Bacteroidales and is defined by several unique features including a predicted tyrosine recombinase, an array of co-oriented short genes (∼100-300 codons) encoding predicted orphan immunity proteins, and conserved intergenic sequence motifs. The rAID system is highly abundant in human gut metagenomes and rAID genes are expressed at high levels in human gut metatranscriptomic datasets, suggesting that it is an important ecological force in the gut [16]. Consistent with this idea, heterologous expression of some rAID genes allows *E. coli* cells to resist intoxication by GA1 and GA2 effectors, and the *orf1* gene of the *B. fragilis* 9343 rAID system confers protection against intoxication during competitive co-culture with GA1-encoding bacteria.

One peculiarity of rAID systems is that the orphan immunity genes found within them typically exhibit limited sequence identity to cognate immunity genes that are genetically linked to cognate effectors, ranging as low as 30% amino acid identity in some cases [16]. This is unusual because it is postulated based on studies examining the structure, function, and molecular mechanisms of T6SS toxin-immunity interactions that immunity genes experience limited sequence evolution, potentially due to the physical interaction required for inhibition of toxin activity by the cognate immunity factor resembling a highly specific lock and key occlusion of an enzymatic active site [6, 21]. Therefore, the question arises whether divergent rAID orphan immunity genes retain the capability to neutralize effector toxicity through protein-protein interaction and, if so, how?

In this study, we explore the structure and function of a broadly distributed class of rAID orphan immunity genes using bacterial growth inhibition assays and biophysical methods combined with metagenomic analyses. In doing so, we uncover evidence that despite extensive sequence divergence, rAID orphan immunity genes encode proteins capable of binding and neutralizing a GA1 effector in a manner that is indistinguishable from that of its cognate immunity protein. We additionally find metagenomic evidence for positive selection and rapid evolution of the effector-binding interface in rAID-encoded orphan immunity genes. Together, our findings reveal new mechanistic insight into the evolutionary and ecological dynamics of interbacterial antagonism among gut bacteria.

## RESULTS

### A widespread immunity gene family is found in both T6SS effector–immunity bicistrons and rAID gene clusters

Previously, we and others identified variable regions in the GA1 T6SS locus of diverse strains of Bacteroidales, a prominent symbiotic taxon in the human gut [10, 12–14]. These variable regions harbor predicted T6SS effector-immunity pairs possessing known features of interbacterial antagonism pathways, including conserved toxin and immunity domains and structural features of toxins such as RHS repeats [20]. One such effector-immunity pair, which we termed E2–I2 based on our prior cataloging of Bacteroidales effector-immunity genes (reference genes from *Parabacteroides* sp. D25, locus tags HMPREF0999_RS10520 and HMPREF0999_RS10515), possesses a conserved WHH nuclease domain and a dual-SUKH immunity domain architecture, respectively (Figure 1A-C) [12, 20, 22]. Among immunity proteins, SUKH (<u>S</u>yd, <u>U</u>S22, <u>K</u>nr4 <u>h</u>omology) domain family members are unique in that they are genetically linked to a diversity of toxins suggesting that their mechanism of toxin inhibition is broadly acting [20, 22]. However, to date, no mechanistic studies have investigated the function of SUKH domains in mediating effector neutralization in the context of interbacterial antagonism.

**Figure 1.**
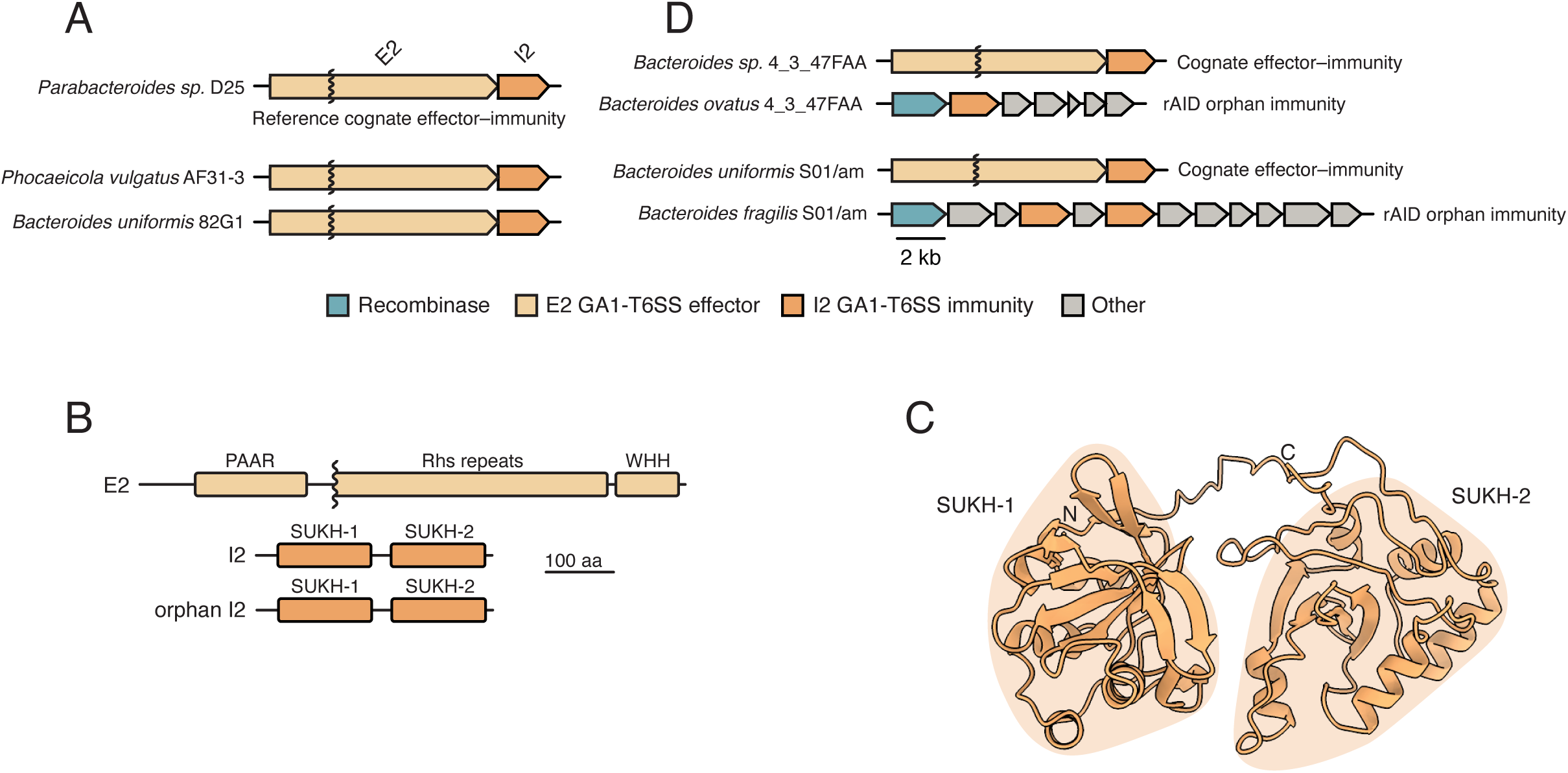
Bacteroidales genomes harbor T6SS and rAID systems containing homologous cognate and orphan I2 immunity genes. **A)** Examples of representative GA1 T6SS E2 effectors and cognate I2 immunity gene loci from indicated reference genomes of Bacteroidales. **B)** Domain diagrams of representative GA1 E2 and I2 proteins. **C)** AlphaFold3-predicted ribbon diagram of the GA1 I2 protein structure highlighting its dual SUKH domain architecture. **D)** Examples of representative cognate GA1 E-I2 loci and orphan I2 loci from strains isolated from the same fecal sample.

To begin to investigate T6SS-associated SUKH immunity factors in human gut bacteria, we downloaded all available Bacteroidales genomes from RefSeq and searched using sequence homology for the presence of E2-I2 bicistrons in GA1 loci, detecting them in 93 genomes from the genera *Bacteroides*, *Phocaeicola*, *Parabacteroides*, and *Alistipes* (Supplemental Table 1). We also detected many dual SUKH-containing genes in 525 genomes with relatively limited homology to cognate I2 (∼30% amino acid identity) encoded within rAID systems (Supplemental Figure 1, Supplemental Table 1). rAID systems encode diverse functional T6SS orphan immunity genes with sequence identity to cognate T6SS immunity ranging from 35-62% [16]. Given that rAID systems are highly prevalent and abundant in adult human fecal metagenomes, we next sought to determine the prevalence of orphan I2 genes in these natural contexts. We used the cognate I2 nucleotide sequence to query the Integrated Gene Catalog (IGC), a database of millions of unique reference genes associated with the human gut microbiome [23]. We found 14 genes with significant shared sequence identity with cognate I2 in the IGC (Supplemental Table 2). We used these genes to query Bacteroidales reference genomes, detecting many SUKH homologs (Supplemental Table 3). Notably, these 101 orphan I2 homologs were identified in only two distinct genomic contexts: within a clearly definable rAID system or in poorly assembled genomes on short scaffolds with features of rAID systems but lacking a contiguously encoded rAID recombinase [16]. Together, our genomic analyses indicate that Bacteroidales genomes harbor abundant SUKH domain-containing orphan I2 genes within rAID systems leading us to hypothesize that they play an important functional role in the ecology of the human gut microbiota.

### Strains encoding orphan I2 genes co-occur in microbiomes with strains harboring cognate E2-I2

During the course of our analysis of Bacteroidales genomes, we made the observation that an E2-I2-containing GA1+ strain (*Bacteroides sp.* 4_3_47FAA) and a rAID orphan I2-containing strain (*B. ovatus* 4_3_47FAA) were isolated from the same adult human donor [24] (Dr. Emma Allen-Vercoe, personal communication). To further investigate the potential for co-occurrence of isolates encoding cognate E2-I2 and orphan I2 genes from other individual microbiomes, we interrogated a large set of Bacteroidales genomes generated as part of a densely-sampled longitudinal fecal collection from 11 adults [25, 26]. Notably, while we identified cognate I2 genes in 7 of 11 individuals in a variety of species in this dataset (Supplemental Table 4), we only detected both orphan I2 and cognate E2-I2 in separate isolate genomes from a single individual, designated S01-am. Examination of the *B. fragilis* S01-am rAID system reveals that it harbors two divergent orphan I2 homologs encoded by open reading frames (ORFs) 3 and 5 of the gene cluster (Figure 1D). Together, these results demonstrate that bacteria harboring rAID-associated orphan immunity genes that are predicted to protect against intoxication by cognate effectors can be co-resident within the same microbiome, suggesting that they confer a fitness advantage to the bacteria that harbor them.

### Divergent I2 proteins protect against E2-mediated growth inhibition

Our genomic studies predict that orphan I2 provides a fitness benefit to bacteria that encode it due its potential capacity to neutralize E2 toxicity. However, many of the orphan I2 genes that we identified in assembled genomes and metagenomic data possess limited sequence identity with cognate I2, in some cases sharing less than 35% identity, which approaches the limit of functional conservation [27]. Therefore, we set out to test if divergent orphan I2 genes are functional and serve to protect bacteria from intoxication by E2. Towards this end, we first utilized a two-plasmid co-expression system in *E. coli* and assessed the viability of cells expressing the predicted nuclease toxin domain (E2_tox_: 149 C-terminal amino acids) and various I2 homologs of increasing sequence divergence. As predicted, E2_tox_ from the reference *Parabacteroides* sp. D25 strain inhibits the growth of *E. coli*. Moreover, growth is restored by co-expressing cognate I2 but not the unrelated I5 immunity protein from the GA1 locus of *B. fragilis* YCH46 (Figure 2A). Interestingly, co-expression of two divergent cognate I2 proteins from *B. uniformis* 82G1 (I2^BU^) and *P. vulgatus* AF31-3 (I2^PV^), each possessing ∼70% amino acid identity to the reference I2, also restored *E. coli* growth under conditions of E2_tox_ intoxication (Figure 2B). More remarkably, co-expression of the *orf3* and *orf5* orphan I2 genes from *B. fragilis* S01-am, which share only 37.7% and 39% sequence identity with cognate I2, also rescued the growth of *E. coli* (Fig. 2B). Finally, expression of a distantly related and divergent (30% amino acid identity) immunity protein possessing only a single SUKH domain from the environmental bacterium *Methylomonas* sp. LW13 (I2^Meth^) failed to restore growth of E2_tox_-intoxicated *E. coli*.

**Figure 2.**
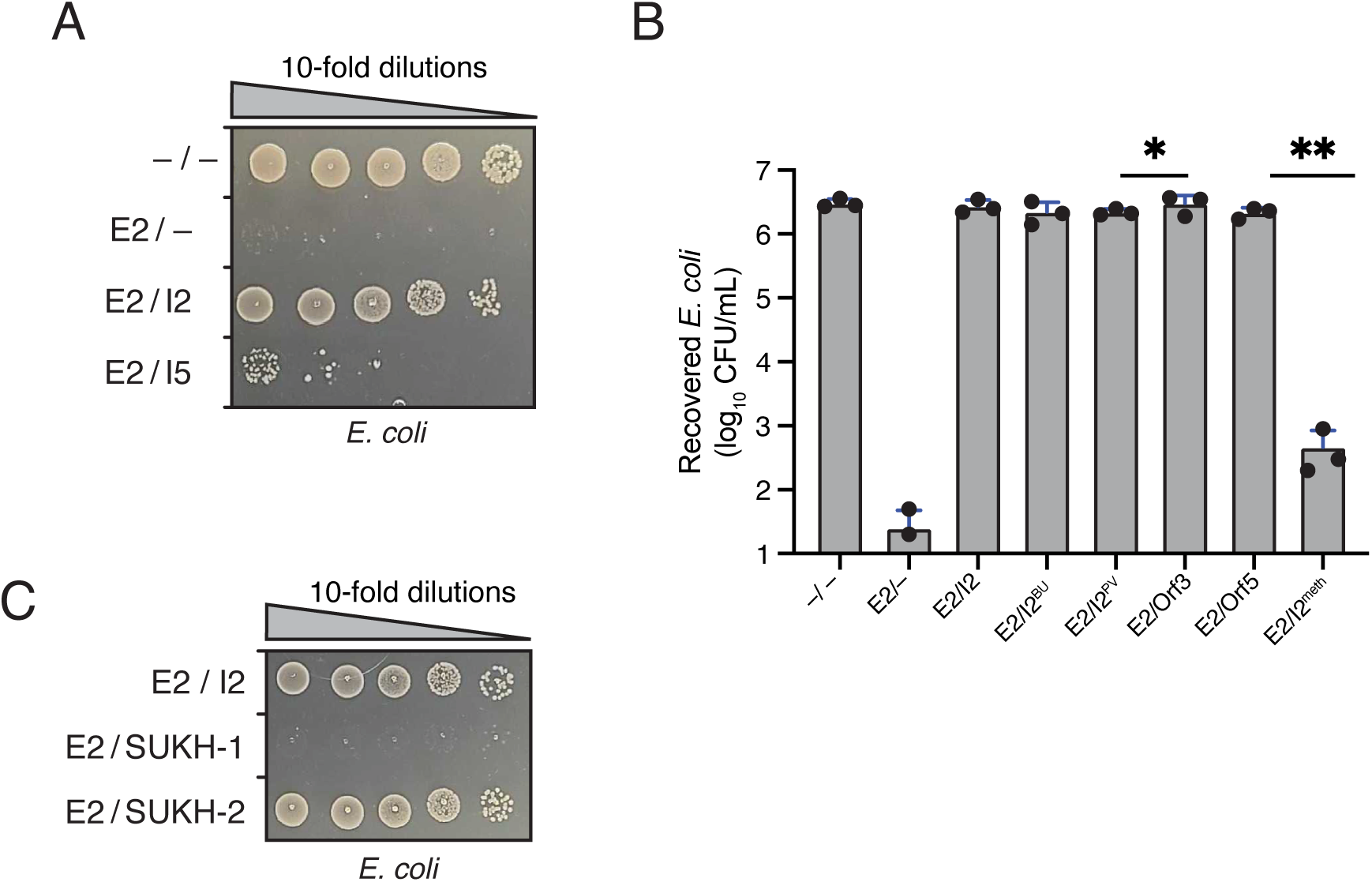
Orphan I2 immunity genes confer protection against intoxication by cognate E2 effectors. **A)** Growth of *E. coli* expressing the indicated E2 effector and immunity proteins (cognate I2, non-cognate I5) on LB plates containing inducers. (–) indicates empty vector controls. **B)** Growth of *E. coli* expressing the indicated effector, cognate immunity, and orphan immunity proteins on LB plates containing inducers. (–) indicates empty vector controls. BU, *B. uniformis*; PV, *P. vulgatus*. **C)** Growth of *E. coli* expressing E2 effector and either full-length I2 or its individual SUKH domains on LB plates containing respective inducers. (–) indicates empty vector controls.

To gain higher resolution insight into how diverse I2 homologs protect against E2 toxicity, we next performed a multiple sequence alignment (MSA) of the experimentally assessed cognate and orphan immunity proteins, including non-protective I2^Meth^. This alignment revealed that both of the SUKH domains found in cognate and orphan I2 proteins share conserved residues predicted to be important for SUKH-specific secondary structure elements (Figure 2C, Supplemental Figure 2A) [22]. However, these positions are also shared by the single SUKH domain of I2^Meth^ and are therefore unlikely to be directly involved in inhibiting the toxicity of E2. We also noted that the overall degree of sequence conservation is higher for the C-terminal SUKH-2 domain than for the N-terminal SUKH-1 domain across the three cognate and two orphan I2 sequences examined, with 46% identity between the SUKH-2 domains of Orf3 and I2, compared to 37% identity for their SUKH-1 domains (Figure 2C, Supplemental Figure 2B). To investigate the relative importance of the two SUKH domains in protecting against E2 intoxication, we performed similar growth assays in *E. coli* in which we co-expressed reference E2_tox_ with truncated cognate I2 fragments consisting of the N- or C-terminal SUKH domains in isolation. In doing so, we found that SUKH-2 is both necessary and sufficient to protect *E. coli* from E2-based intoxication (Figure 2C). Collectively, these data indicate that Bacteroidales orphan I2 genes encode functional immunity proteins that can protect bacteria from intoxication by E2 and that a single SUKH domain is sufficient for this function.

Based on our *E. coli* data, which support the capacity of orphan I2 to neutralize E2-mediated toxicity, we hypothesized that endogenous rAID-encoded orphan I2 genes harbored by Bacteroidales species could also perform this function *in vivo*. To explore this possibility, we turned to the genetically tractable laboratory strain *B. fragilis* NCTC 9343, which possesses a rAID system harboring two orphan I2 homologs at ORFs 5 and 10 (referred to *orf5* and *orf10*, Figure 3A), which encode proteins with 37.9% and 36.1% amino acid identity to cognate I2, respectively. We generated isogenic in-frame chromosomal deletions of each gene separately and together in the same strain. In each of the resulting mutant strains and in the wild-type strain, we integrated an anhydrotetracycline (aTc)-inducible E2 expression construct at the *att1* chromosomal site. In WT or in *Δorf5* cells, induction of E2 expression did not result in a statistically significant difference in recovery of intoxicated *B. fragilis* cells compared to uninduced cells (Figure 3B). By contrast, the viability o f a strain lacking *orf10* was significantly reduced under inducing conditions and this reduction in viability was exacerbated in a strain lacking both *orf5* and *orf10*, which exhibited 1000-fold reduction in viable cells recovered. Together, these data indicate that the two orphan I2 genes in the *B. fragilis* 9343 rAID system provide protection against E2 intoxication *in vivo* and that gene dosage likely underlies their additive protective effect. We further sought to assess if orphan I2 genes harbored by strains isolated from E2-containing microbiomes were also capable of protecting cells from E2 intoxication. Towards this end, we integrated the same aTc-inducible E2 expression construct into *B. ovatus* 4_3_47-FAA and *B. fragilis* S01-am, which possess one and two copies of orphan I2, respectively (Figure 1D). We found no difference in viable cell count between induced and uninduced conditions, indicating that these strains are protected from E2 intoxication (Figure 3C).

**Figure 3.**
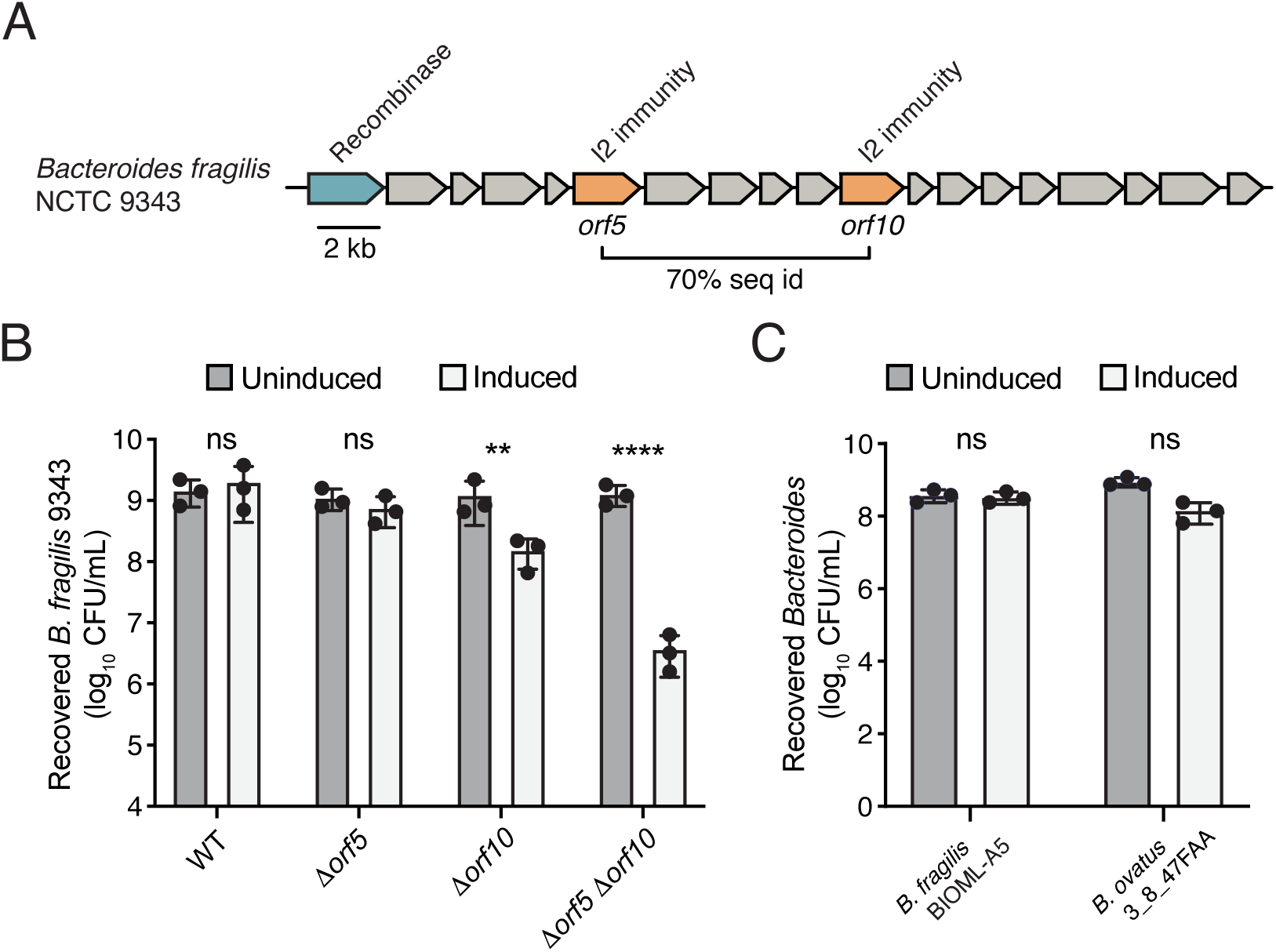
Orphan I2 immunity genes protect Bacteroidales against intoxication by cognate E2 effectors. **A)** The *B. fragilis* NCTC 9343 rAID system. Predicted tyrosine recombinase (blue) and orphan I2 genes at open reading frame (ORF) positions 5 and 10 (orange) are indicated. **B)** Growth of WT and genetically manipulated *B. fragilis* NCTC 9343 strains on BHIS plates containing or lacking inducer (anhydrotetracycline, aTC) for expression of E2, quantified by colony forming units (CFU) depicted on a log10 scale. **C)** Growth of *B. fragilis* S01-am (BIOML-A5) and *B. ovatus* 3_8_47FAA on BHIS plates containing or lacking aTC for expression of E2. ** p <0.001; *** p<0.0001; ns, not significant.

### Cognate and rAID encoded I2 proteins bind E2 with nanomolar affinity

T6SS immunity proteins typically inhibit cognate effector proteins via the steric occlusion of enzymatic active sites and as a result of the highly specific protein-protein interaction required for this type of binding to occur, they exhibit limited sequence divergence across homologs presumably because of purifying selection to maintain the binding interface [6, 21, 28, 29]. For example, highly conserved type VI secretion immunity proteins in *Proteus mirabilis* (e.g. 89.6% amino acid identity identity) do not confer protection against non-cognate toxins [30]. Nonetheless, our growth inhibition experiments in both *E. coli* and *B. fragilis* indicate that divergent I2 homologs possess more broadly acting inhibitory properties. To test if this is due to a high affinity protein-protein interaction that is maintained across diverse I2 homologs, we next set out to measure the binding affinity between E2_tox_ and various I2’s using purified proteins. To overcome the extreme toxicity of E2_tox_ towards *E. coli*, we first expressed and purified his_6_-tagged E2_tox_ as a 1:1 complex with untagged cognate I2 via nickel affinity and size exclusion chromatography coupled with multi-angle laser light scattering (Figure 4A and 4B). The resulting complex was then denatured to remove unfolded I2 and column-bound E2_tox_ was subjected to on-column refolding in renaturation buffer prior to its elution from the column in its native state (Figure 4C). Cognate I2 and each of the I2 homologs was expressed and purified using standard methodologies.

**Figure 4.**
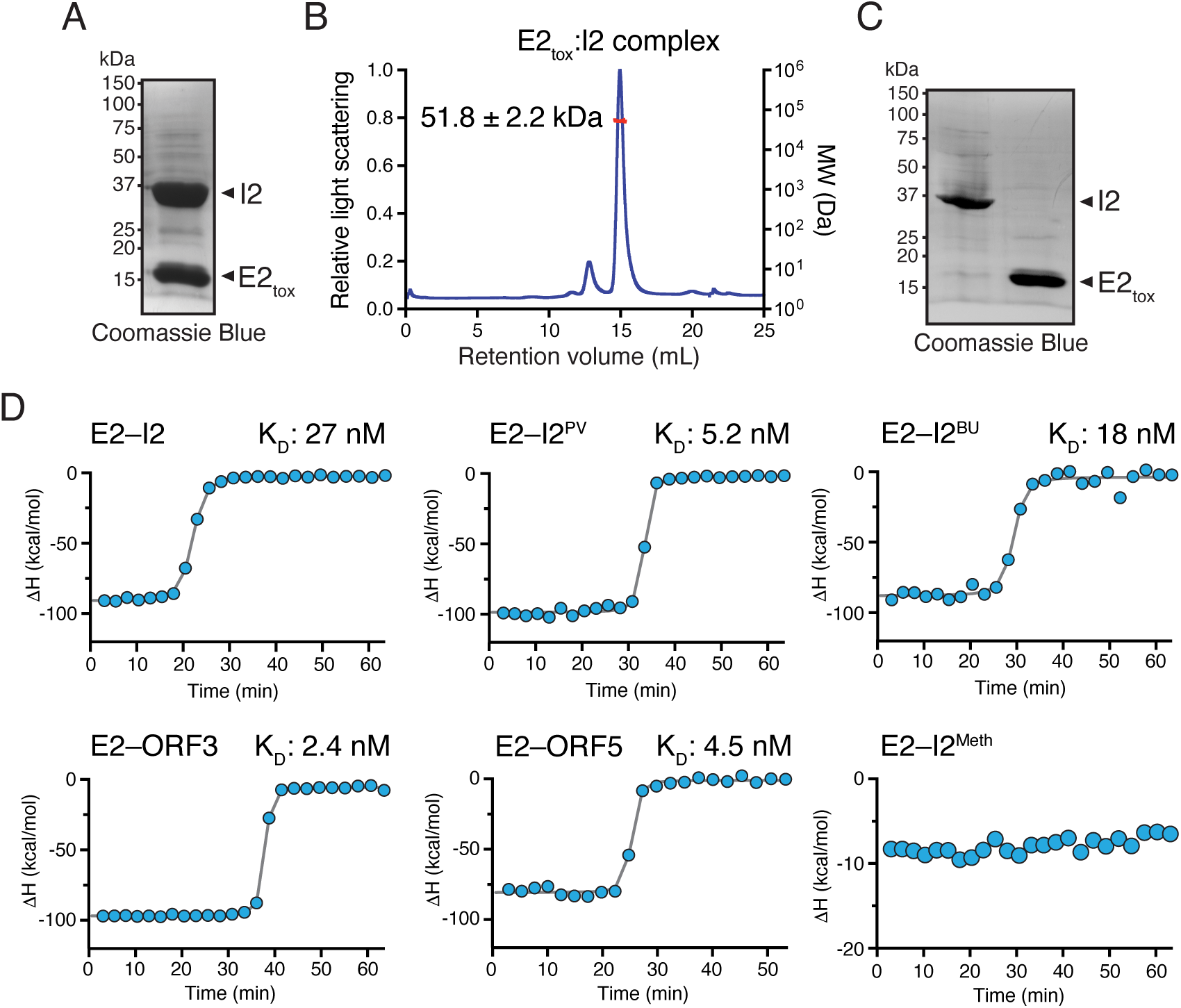
Cognate and rAID-encoded I2 immunity proteins exhibit nanomolar binding affinity to E2 effector. **A)** Coomassie-stained SDS-PAGE gel showing purification of recombinant E2_tox_-I2 complex after co-expression in *E. coli* and co-purification by nickel affinity chromatography. **B)** SEC-MALS chromatogram and molecular weight analysis of E2_tox_-I2 complex. The indicated molecular weight corresponds to a 1:1 complex. **C)** Coomassie-stained SDS-PAGE showing separation of E2_tox_ and I2 after denaturation of the E2_tox_-I2 complex, removal of I2 by denaturing wash, and refolding of E2_tox_. **D)** Quantification of cognate and orphan I2 binding affinity to E2_tox_ by isothermal titration calorimetry. The dissociation constant (K_d_) for each complex is indicated.

We next measured the affinity between purified E2_tox_ and various I2 proteins using isothermal titration calorimetry (ITC). In line with our growth inhibition data, cognate I2 and each of the four microbiome-derived homologues of I2 that protect against E2-mediated toxicity interacted with E2_tox_ with nanomolar affinity (Figure 4D). By contrast, non-protective I2^Meth^ exhibits no detectable binding affinity towards E2.

Collectively, these data demonstrate that despite the differing levels of sequence identity between cognate I2 and the four protective homologs from Bacteroidales, they all possess the ability to form high affinity complexes with E2 and block its ability to inhibit bacterial growth.

### An acidic patch on I2 facilitates its ability to inhibit E2 antibacterial activity

To better understand the molecular basis of inhibition of E2 toxicity by cognate and orphan I2 homologs, we examined our MSA of I2 and five protective non-cognate immunity proteins to find regions of I2 that are the same among the protein sequences that protect against E2_tox_-mediated toxicity but are different from the non-protective I2-Meth homologue (Figure 2C, Supplemental Figure 2). Overall, we found 27 residues that meet that criterion. This list was further refined by examining a high confidence AlphaFold3 model of the E2-I2 complex and eliminating residues that are not located at the interaction interface (Figure 5A). Interestingly, five out of the remaining six residues (E228, D229, D231, E234, and N236) possess electronegative side chains suggesting that negative charge is a key feature of I2 proteins that can inhibit E2 activity. To test this prediction, each residue was mutated to either arginine or lysine to assess the importance of charge for immunity function. Perhaps surprisingly, given the drastic nature of charge-swapping mutations, combinations of single- and double-point mutants of I2 could still protect against E2-mediated toxicity. It was not until we constructed a quintuple variant of I2 (I2^5x^) was the ability to protect against E2 lost (Figure 5B). Importantly, western blot analysis indicates I2^5x^ expression levels are equivalent to that of the wild-type protein indicating that these mutations unlikely impact the stability of the mutated protein (Supplemental Figure 3). Overall, these results support our hypothesis that the acidic residues located on I2 play a critical role in conferring binding specificity to E2. Additionally, the mutational tolerance observed during these studies indicates that this binding interface can be maintained even as the orphan I2 sequence diverges from cognate I2.

**Figure 5.**
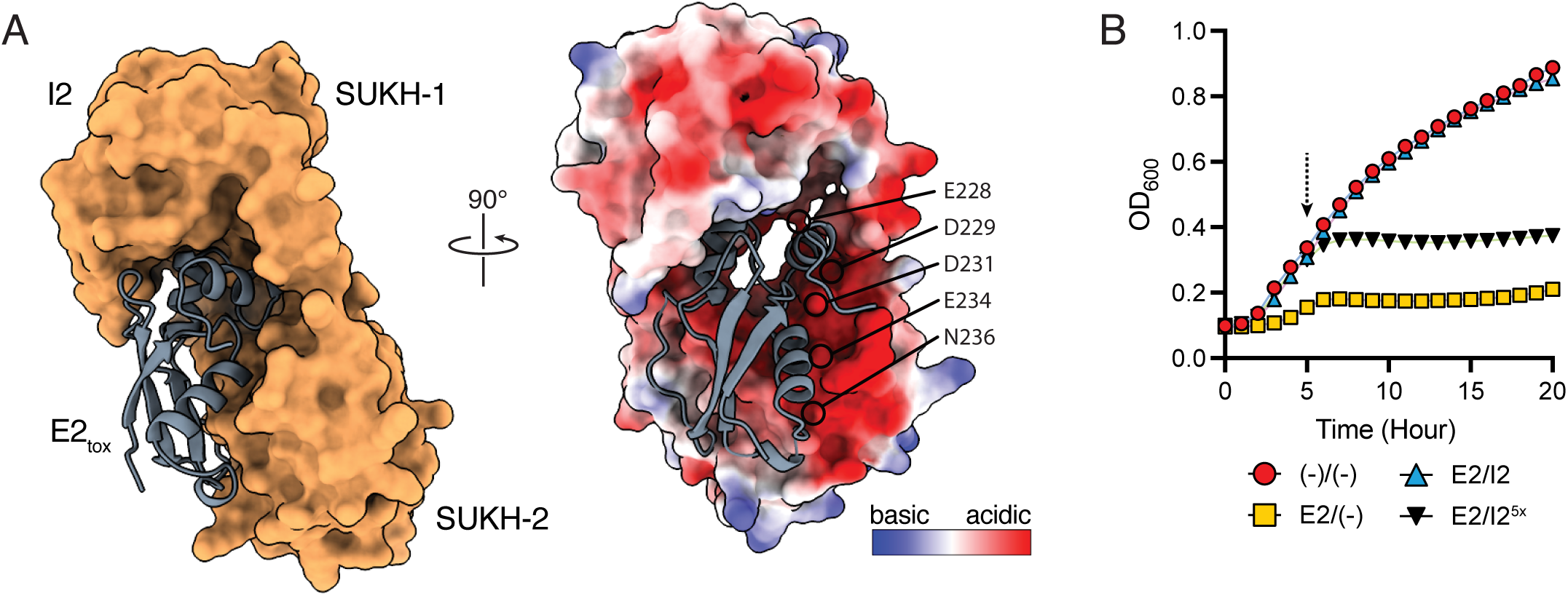
Alphafold3 modeling of the E2-I2 complex reveals an interaction interface that is governed by electrostatic interactions. **A)** Overall model of the complex and identification of the E2-I2 interaction interface. The complex is modelled with an iPTM score of 0.95, a predicted buried surface area of 2314.2 Å^2^, and a predicted free energy of binding (ΔG) of –13.9 kcal/mol. The predicted electrostatic surface potential of I2 (right) contains a patch of negative charge potential around the E2 interface comprised of several acidic residues (E228, D229, D231, E234, N236). **B)** The identified acidic residues are required for E2 toxin neutralization. Growth curves of *E. coli* harboring empty expression plasmids (–), or plasmids for expression of effector (E2), immunity (I2), or an immunity variant with Arg/Lys substitutions of the five acidic residues shown in A (I2^5x^).

### Analysis of human gut metagenomes reveals orphan I2 is associated with an E2-dependent increase in abundance

Thus far, our genomic and functional studies indicate that rAID-encoded I2 proteins protect bacterial cells from E2 intoxication through maintenance of a tight protein-protein interaction despite significant sequence divergence from cognate I2. This implies that in the context of the gut microbiota, bacteria that possess I2-encoding rAID systems would receive a fitness benefit, but likely only under conditions in which E2 exerts significant selective pressure. Evidence to support this conjecture could come from the analysis of metagenomic data and we therefore examined the distribution and abundance of cognate E2-I2 and orphan I2 genes across adult metagenomic samples from the Human Microbiome Project [24]. We found that cognate E2 and I2 genes were detected in only 55 and 56 out of 469 microbiomes, respectively, and were co-detected in the same microbiome in all but one case (Figure 6A). By contrast, orphan I2 genes were found in 413 of 469 samples. Despite the wide prevalence of orphan I2 across metagenomes, we found relatively sparse patterns of co-detection with cognate E2, providing an opportunity to assess if co-detection is associated with differences in abundance. We therefore quantified orphan I2 gene abundance in those metagenomes in which we detected E2 and compared those abundances to that of metagenomes in which E2 was absent. Remarkably, we found that E2-positive metagenomes exhibit significantly higher orphan I2 gene abundances than those without E2 (Figure 6B). This observation provides correlative evidence that orphan I2 confers a fitness advantage to bacteria in microbiomes in which T6SS-dependent E2 intoxication is exerting a selective pressure on neighboring Bacteroidales strains.

**Figure 6.**
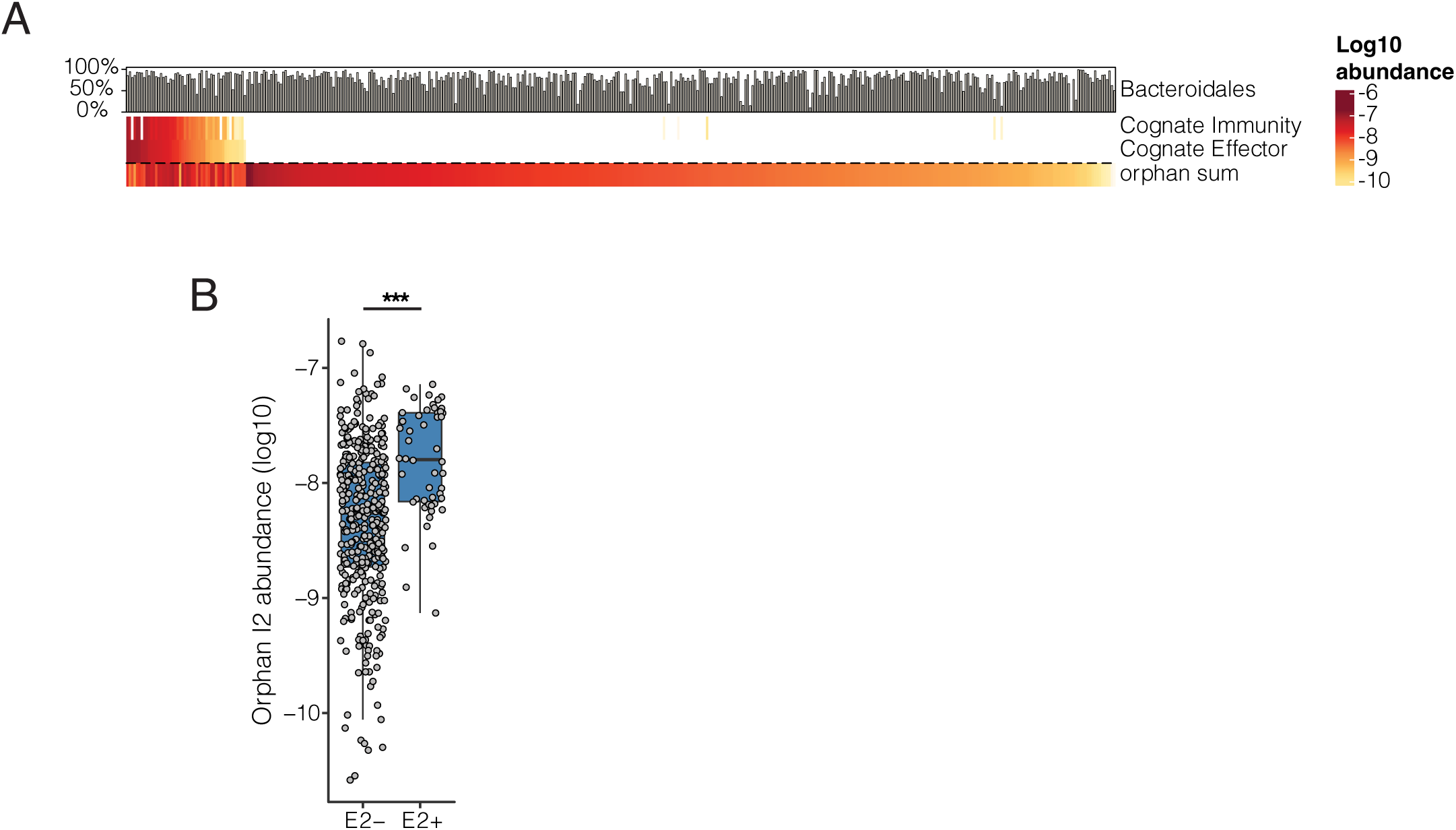
Orphan I2 immunity genes are more abundant in metagenomes that harbor cognate effectors than in those that do not. **(A)** Heatmap depicting quantification of abundance (log_10_ scale) of cognate effector, cognate immunity, and orphan immunity genes (rows) in adult human fecal metagenomic samples (columns) from the HMP. Bar plots indicate the abundance of the order Bacteroidales in each sample **(B)** Quantification of orphan I2 gene abundance in adult human fecal metagenomes in which cognate E2 was either detected or absent. P < 0.00001, Wilcoxon rank sum test, N = 469.

### SNP analysis of genomes and metagenomes from S01-am

Our metagenomic survey suggests that orphan I2 genes provide a fitness benefit to Bacteroidales strains inhabiting E2-containing microbiomes. This finding, coupled with the high level of sequence divergence we observed between orphan and cognate I2 genes, led us to hypothesize that orphan I2 may evolve under elevated rates of non-synonymous substitutions reflective of positive selection. While assessment of signals of positive selection in specific genes across independent bacterial lineages can be challenging due to the rapid rate of mutation in bacterial genomes, recent efforts have dramatically advanced the ability to conduct such analyses in the context of the gut microbiome [26, 31, 32]. We sought to leverage longitudinal evolutionary dynamics within the same bacterial lineages using metagenomic data from human subject S01-am to assess if orphan immunity genes were subject to positive selection [25, 26]. We first examined microbiome dynamics in the S01-am samples and detected a significant shift in the species-level composition of this community over time (Figure 7A, PERMANOVA, p < 0.001). Mirroring this shift in species composition, the abundances of both cognate E2-I2 and orphan I2 genes also changed, with the cognate E2-I2 genes generally increasing over time whereas orphan I2 genes exhibit an oscillating pattern of abundance (Figure 7B). To determine which species possess these respective genes, we next queried isolate genomes from S01-am. We detected cognate E2-I2 genes primarily in isolates of *Phoecicola vulgatus* and a single isolate of *B. uniformis*. Detection of the same T6SS genes in different species is expected and is consistent with the previously observed intra-microbiome mobilization of the GA1 T6SS between Bacteroidales species [10–12, 26, 33]. By contrast, we found orphan I2 genes in several species in S01-am, including *B. fragilis*, *P. vulgatus, B. ovatus,* and *Alistipes onderdonkii*. We confirmed that the orphan I2 genes encoded in *P. vulgatus* isolate genomes were observable in metagenomes by verifying that there is a correlation between *P. vulgatus* and *orf3*/*orf5* genes (r=0.74 for both), and we observe a similar effect for the cognate E2 and I2 genes encoded in *B. uniformis* (r=0.77 for both in the metagenome data). By contrast, there was no significant correlation for *B. fragilis*. Examining the abundance of these species over time showed that they oscillated in a manner similar to the genes that they likely encode. By contrast, there was no significant correlation between the metagenome abundance of *B. fragilis* and *B. ovatus* at the metagenome level and only a very weak correlation for *B. salyersiae* (r=0.23 for E2, r=0.22 for I2, both p<0.01).

**Figure 7.**
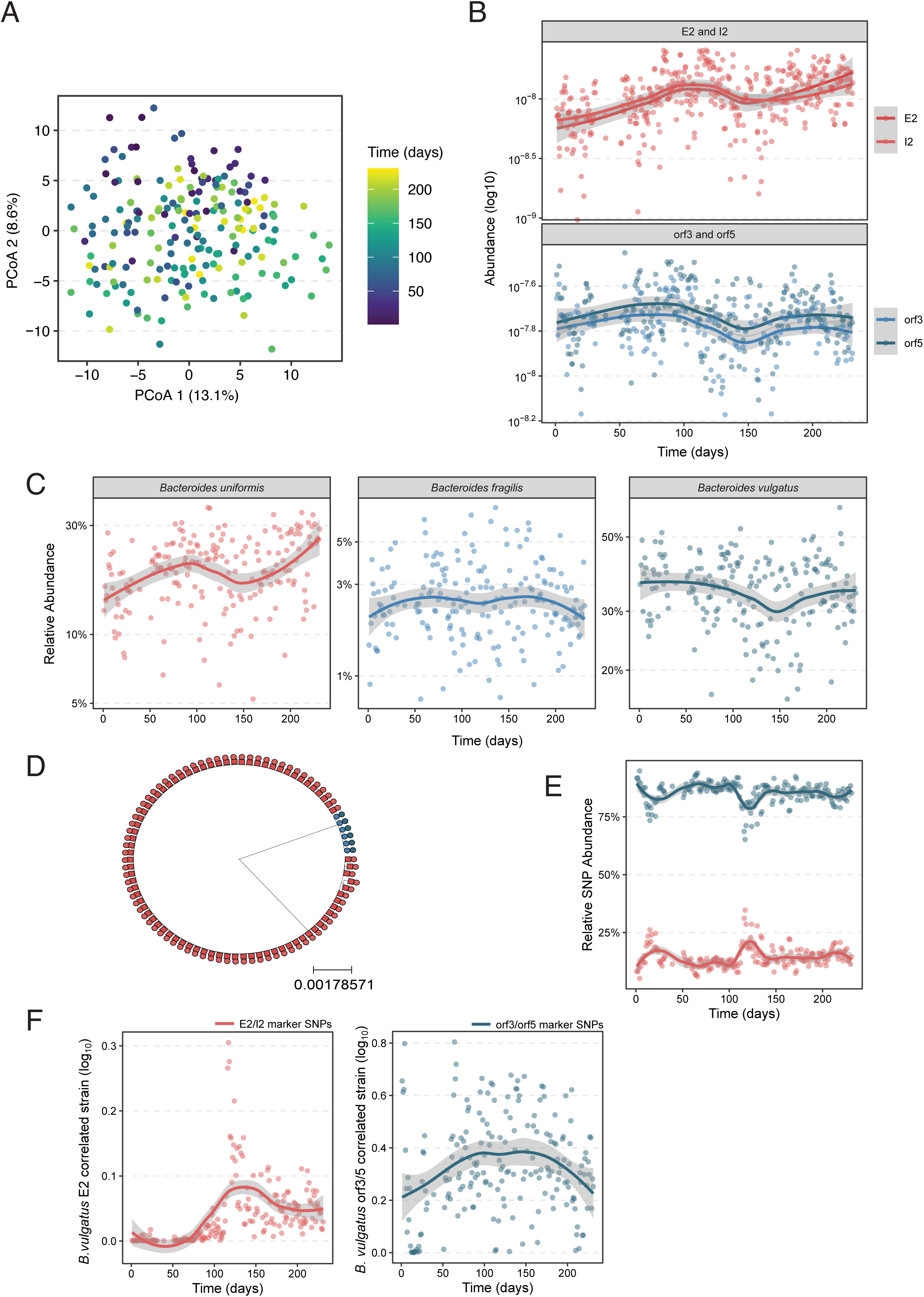
The S01-am microbiome exhibits complex population dynamics. **A)** Principal component plot showing significant shift in microbiota composition for individual S01-am over the collection time series (PERMANOVA p<0.001). Points indicate individual samples colored by time at collection. **B)** Gene abundance (log_10_) plots for cognate E2 and I2 (left panel) and orf3/5 in S01-am over time. Note that the y-axis scales differ between panels. **C)** Relative abundance plots over time in S01-am metagenomic samples for indicated species. **D)** Phylogenetic tree depicting *P. vulgatus* strains isolated from S01-am samples annotated by E2-I2 (red) or orphan I2 (blue). **E)** Relative abundance plotted over time for *P. vulgatus* strain-specific marker SNPs. **F)** Abundance of the *P. vulgatus* E2-correlated (left panel, red) and orphan I2-correlated strains in S01-am metagenomic samples.

We were particularly interested in *P. vulgatus* because it was surprising that isolates of this species encoded either orphan immunity or cognate effector-immunity genes but not both. While our previous examination of NCBI reference genomes did not identify any instances these genes being present in the same genome, multiple strains of *P. vulgatus* have been reported to co-colonize the human gut [34] and therefore we hypothesized that in the S01-am microbiome, distinct strains harbor cognate and orphan genes. To investigate this possibility, we constructed a phylogenetic tree of all *P. vulgatus* isolate genomes from S01-am samples using species-specific marker genes. This analysis revealed at least two phylogenetically distinct *P. vulgatus* strains with a higher number of isolate genomes contain E2 compared to those containing orphan I2 (Figure 7D). We then used our phylogenetic tree to identify strain-diagnostic single nucleotide polymorphisms (SNPs) that would allow for the precise quantification of these *P. vulgatus* strains in S01-am metagenomes. In contrast to what we expected from our isolate genome analysis (Figure 7D), strain-specific SNPs associated with the E2 strain were significantly lower in relative abundance than those from the orphan I2 strain over the entire longitudinal dataset (Figure 7E), implying that isolation recovery of each strain by culturing was disproportional with respect to the true *in vivo* population proportions as detected by metagenomic sequencing analysis. At approximately the 300-day timepoint, we observed a significant deviation in SNP relative abundances associated with an expansion of the E2-containing *P. vulgatus* strain over time and a corresponding decrease in the *P. vulgatus* orphan I2 strain.

To confirm that these changes in SNP abundance corresponded to distinct strains in the metagenome, we used StrainFacts to identify *P. vulgatus* strains in the S01-am microbiome [35]. There is no simple empirical way to determine the true number of strains for a given species in a metagenome via StrainFacts, but we were interested in strains that represent the cognate and orphan encoding subpopulations that we inferred exist from S01-am isolate genomes. Therefore, we used the detection of metagenomic strains that had a reasonable correlation to cognate and effector immunity genes to help estimate strain number (see Methods). With 4 strains, we found one strain that was correlated to the cognate effector (r=0.43) and another strain that was correlated to orphan *orf5* (r=0.45) and to a lesser extent orphan *orf3* (r=0.22). Quantitation of changes in relative abundance over time for these strains exhibited dynamics similar to that of the cognate and orphan genes (Figure 7F). Remarkably, the E2-associated strain appeared to be at very low abundance before blooming after day 200, a pattern not observable when only looking at the genome derived SNPs. To confirm that this pattern of increased relative abundance was observable at the gene level and not just the SNP level, we identified genes found in the E2/I2 encoding genomes, quantified their abundance in metagenomes, and found that many of those genes exhibited similar dynamics (Methods, Supplemental Figure 4).

The above genomic and metagenomic analyses reveal intriguing population dynamics in S01-am, including the dramatic expansion of an E2-containing *P. vulgatus* subpopulation. We hypothesized that this expansion might result in increased T6SS-mediated selective pressure on orphan I2 genes encoded by co-resident species in S01-am since numerous Bacteroidales species encode homologs of these genes. To investigate this idea, we sought to understand gene-level population dynamics across S01-am longitudinal samples. Although cognate effector-immunity genes did not display any evidence of SNPs, we found numerous non-synonymous SNPs in the *orf3* gene that clustered into two distinct regions of the protein corresponding to the two SUKH domains (Figure 8A). These SNPs associate between species as is expected given that they were identified from Bacteroidales genomes, but we also identify segregation within species with *B. fragilis* and *P. vulgatus* being especially noteworthy (Figure 8B). Tightly clustered SNPs are evidence of selection because neutral evolution should occur uniformly. To gather more evidence, we assessed if the abundance of these variants changed over time. For this purpose, we treated blocks of adjacent SNPs co-detected in the same genome as a single genotype and quantified them together. Remarkably, some SNPs in orphan I2 genes exhibited significantly altered abundance over the S01-am sampling period (540 days, Figure 8C). In particular, a non-synonymous SNP (Y->F at position 223) showed the clearest evidence of increasing over time and it segregates within strains of *B. fragilis* and *A. onderdonkii*. We also examined SNPs in *orf5* but found only two that were found in a region that was identical to *orf3* and thus can only be distinguished at the genome level and not the metagenome level. A third SNP whose metagenome abundance was extremely low was also identified (1 count in each of 5 samples) and therefore no evidence of selection was possible.

**Figure 8.**
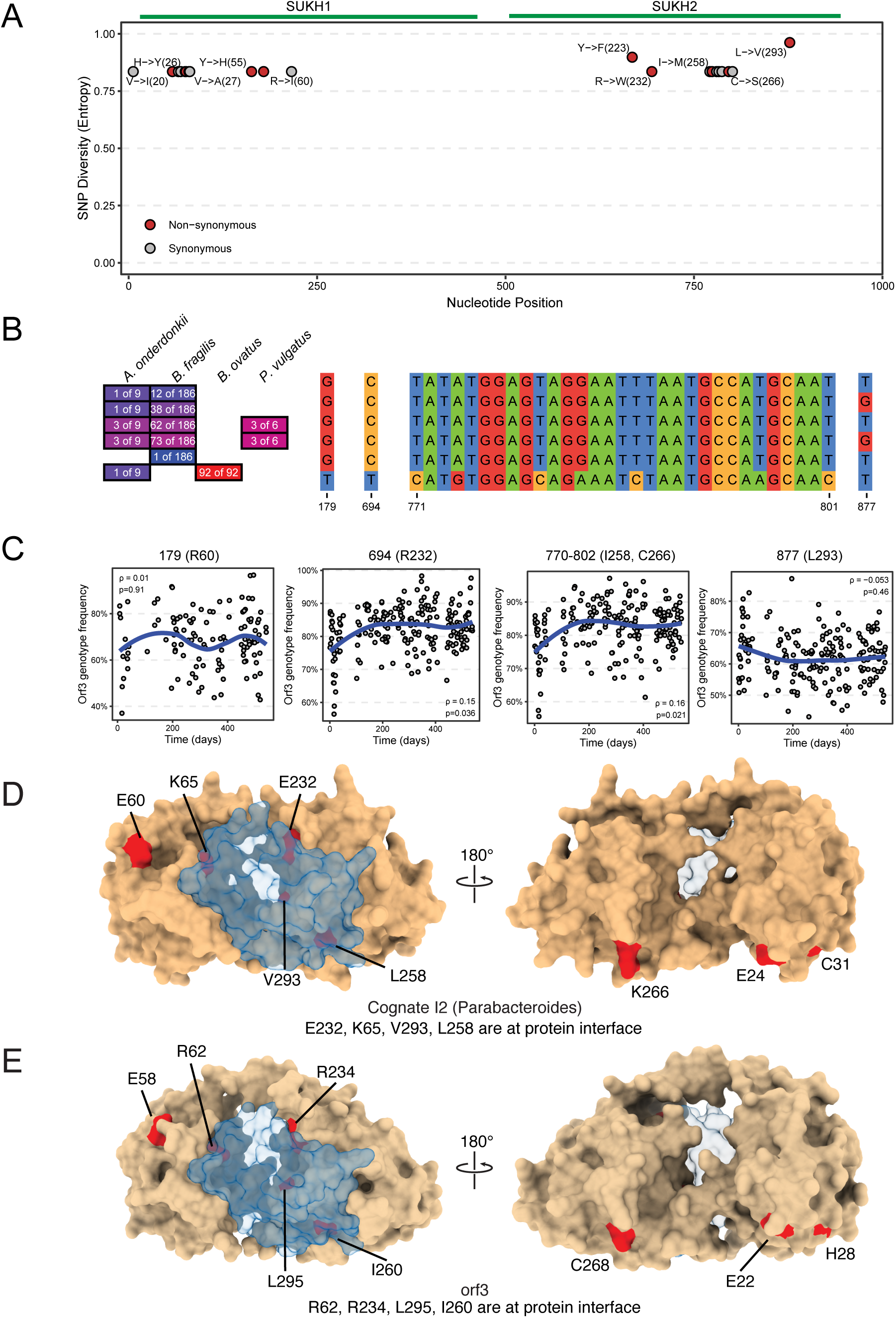
Rapid within-person evolution of orphan I2 sculpts the E2-I2 interaction interface. **A)** SNP diversity detected in the S01-am metagenomes at each position in the orphan I2 gene (Y-axis) plotted against nucleotide position (X-axis), non-synonymous (red), synonymous (gray). Amino acid positions are indicated next to select SNPs. **B)** Quantification of SNP genotypes from S01-am isolate genomes of the indicated species. Nucleotide alignment displays blocks of shared sequence across strains and species for positions of interest. **C)** Scatter plots depict quantification of three SNPs determined to be significantly changing in abundance over time in the S01-am microbiome. **D)** AlphaFold3 predicted structure of the E2 (transparent) and cognate I2 (tan) complex. Non-synonymous mutations are highlighted in red. **E)** AlphaFold3 predicted structure of the E2 (transparent) and orphan I2 (tan) complex. Non-synonymous mutations are highlighted in red.

Last, we sought to understand the potential functional consequences of the rapidly evolving non-synonymous SNPs we identified. Mapping these SNPs onto Alphafold3-predicted I2 structures revealed their enrichment at the predicted E2 protein-protein interaction interface for both cognate and orphan I2, including several acidic residues that could contribute to electrostatic interactions (red highlights in space-filling model, Figure 8D-E). Contrary to the null expectation of a random distribution of these SNPs throughout the protein, their clustering at the potential interface underscores that their positive selection might drive functional changes beneficial in neutralizing E2 toxicity.

## DISCUSSION

### Summary of findings

The study of bacterial defense systems is a rapidly developing field with many notable advances being made in the discovery and characterization of antiphage defense systems [36]. However, significantly less is known about other types of defensive mechanisms such as those that provide bacteria with protection from interbacterial antagonism [18, 19]. Here we have investigated the function, evolution, and ecological dynamics of the I2 family of SUKH-domain-containing orphan immunity genes harbored by defensive rAID systems of Bacteroidales. Our studies revealed that these orphan immunity factors, despite significant sequence divergence from their cognate homologs, can protect against E2 effector toxicity. Individual microbiomes with T6SS-associated cognate E2-I2 genes frequently also harbor strains encoding functional orphan I2 immunity genes, and we provided evidence that interactions between the T6SS and orphan immunity harbored by rAID systems are ecologically relevant since they possess the potential to alter the outcome of T6SS-mediated antagonistic interactions. rAID orphan immunity genes confer a selective advantage to bacteria that carry them and this is reflected by elevated rates of non-synonymous mutations in orphan I2 genes in longitudinal microbiome samples, suggesting that orphan I2 genes are the targets of positive selection.

### Implications for rAID system evolution and function

We found that Bacteroidales genomes harboring orphan I2 genes often encode multiple divergent copies within the same rAID system. The results from our *Bacteroides in vitro* assays imply that orphan immunity gene copy number (and by inference protein dosage) is relevant for determining the outcome of effector intoxication since a *B. fragilis* 9343 strain lacking both of its orphan I2 genes exhibits higher susceptibility to E2 toxicity than strains lacking only one I2 gene. Similarly, metagenomes with E2 genes have higher abundances of orphan I2 genes than metagenomes without suggesting that an increase in orphan I2 copy number is beneficial in the microbiome, although we could not determine if this finding reflects higher abundance within or between genomes (e.g. by gene duplication or horizontal gene transfer between different strains or species, respectively) or merely increased population size of the bacteria encoding it. The mechanism by which a particular rAID system might acquire two homologous copies of an orphan immunity gene such as I2 is not yet clear. Possibilities include duplication of a pre-existing rAID gene or multiple independent acquisition events into the same gene cluster via horizontal gene transfer. Distinguishing between these possible scenarios and defining the mechanism of rAID biogenesis is an area of active research likely to yield insight into the evolution of these defensive systems.

### Interbacterial antagonism and orphan immunity dynamics within the gut microbiome

Although GA1-encoded E2 genes are relatively rare, possibly reflective of the broad diversity of the GA1 effector repertoire [10, 12], orphan I2 genes are exceeding common across adult human gut microbiomes. One implication of such a high prevalence is that functional orphan I2 genes may provide a barrier against invasion into an established microbiome by E2-containing bacteria because the fitness benefit of T6SS activity mediated by E2 would be reduced or eliminated. A notable feature of the GA1 T6SS is that it is found on an integrative and conjugative element (ICE) and can therefore be transferred broadly across co-resident strains and species of Bacteroidales within the same microbiome [10, 33]. We find evidence that identical defensive rAID systems are also present within multiple co-resident strains and species in genomes from S01-am, suggesting that they are also mobilizable and that a potential fitness benefit of an orphan immunity gene like I2 could be rapidly spread within a microbiome. Intriguingly, none of the S01-am genomes we analyzed had both an orphan I2-containing rAID system and an E2-containing GA1 T6SS. This finding suggests the potential for intra-genomic incompatibility between matched rAID orphan immunity and cognate genes, with potential implications for species- and strain-level interactions among Bacteroidales and corresponding co-existence patterns in human gut microbiomes.

### Positive selection at the effector-immunity interface

We used AlphaFold3 to predict the protein-protein interaction surface between E2 and I2. This analysis revealed an interaction interface comprised of extensive electrostatic interactions. Experimental analysis of this interface revealed surprising mutational tolerance as alteration of single amino acids was not sufficient to abolish I2’s ability to protect against E2-mediated toxicity. It was not until multiple mutations to charged residues were introduced into I2 that we were able to observe loss in immunity function.

T6SS immunity proteins are known to bind to and inhibit the toxic activities of their associated effectors through several distinct mechanisms, including electrostatic interactions as observed here. The most common mechanism identified to date involves the physical occlusion of an effector’s active site by the bound immunity protein [28, 37].

As noted by Bosch and colleagues [38], this type of inhibition can occur by an immunity protein preventing an effector’s substrate from accessing its catalytic centre (termed ‘capping’) or by some portion of the immunity protein, typically a loop region, physically occupying the substrate binding site within an effector’s active site (termed ‘plugging’). A more recently identified mechanism of effector inhibition is structural disruption, whereby the binding of the immunity protein induces major structural rearrangements in the effector such that it is no longer in a catalytically competent state [38]. Our biochemical analysis of the I2 family of immunity proteins indicates that their SUKH domains likely inhibit E2 effectors using the former of these two mechanisms. Such a mechanism of inhibition is expected to have greater mutational tolerance because as long as the free energy of complex formation remains favorable, inhibition is likely to still occur. By contrast, structural disruption likely requires a series of stepwise interactions and conformation changes that requires the participation of many specific residues that if mutated would be more likely to disrupt immunity function. Therefore, we expect that immunity proteins with broadly inhibitory activities are more likely to act through a physical occlusion mechanism.

Our analysis from the longitudinal S01-am metagenomes indicates that residues at the predicted effector-orphan immunity interface are the targets of positive selection, implying that there are fitness benefits to mutational alterations at this interface that could potentially include subtle gain of function effects. These results may indicate physiological differences between the delivery of single effector proteins during T6SS-mediated interactions in the gut versus an *in vitro* situation in which both effector and immunity are over-expressed and subtle effects are masked. Since the S01-am metagenomic data we analyzed was composed of short sequencing reads, our analysis did not allow us to distinguish if the detected non-synonymous SNPs might reside in the same gene harbored within the same genome (e.g. linked mutations) or represent different alleles present in the same microbiome but encoded by different genomes. Recent work characterizing the GA3-specific effector Bte1 and its associated cognate immunity factor Bti1 identified evidence of co-evolution reminiscent of the findings presented here [39]. Such signatures of rapid evolution are consistent with broad and ongoing evolutionary arms races between T6SS-encoded effectors and orphan immunity genes across human gut Bacteroidales.

## Acknowledgements

We thank members of the Dartmouth College Joint Evolutionary Microbiology Meeting (JEMM) and M2P2 communities for feedback, Dartmouth College Research Computing for the use of computing resources, the Dartmouth BioMT core (P20-GM113132), and the Dartmouth Genomics and Molecular Biology Shared Resource at Dartmouth, which is supported by NCI Cancer Center Support Grant P30CA023108 and NIH S10OD030242 awards. We thank Britt Berdy at MIT for sharing strains from individual S01-am and Emma Allen-Vercoe at University of Guelph for sharing strains from the HMP. We are deeply grateful for generous funding from the National Institutes of Health under grants R00GM129874 and R35GM142685, and start-up funds from the Dartmouth College Geisel School of Medicine to BDR. This project was also supported by a seed grant from the David Braley Centre for Antibiotic Discovery (to JCW). PH was supported by the Dartmouth Host-Microbe Training Grant T32AI007519. JCW is the Canada Research Chair in Molecular Microbiology and holds an Investigators in the Pathogenesis of Infectious Disease Award from the Burroughs Wellcome Fund.

## METHODS

### Cloning and Plasmid Construction

All molecular cloning was performed using standard restriction enzyme- or isothermal assembly-based methodologies (NEB). For toxicity experiments in *E. coli*, toxin genes were cloned into pSCrhaB2 and immunity genes were cloned into pPSV39-CV [40] [41]. pETDuet-1 (Novagen) was used for overexpression of E2-I2 complexes, with effector genes cloned into MCS-1 and immunity genes into MCS-2. pET29b (Novagen) was used for overexpression of immunity proteins in isolation. All Bacteroidetes genes were codon optimized for expression in *E. coli* (Genscript). Site-specific mutants generated for this study were constructed by overlap extension PCR. All plasmids were sequenced by Genewiz Incorporated.

### Toxicity Assays

*E. coli* toxicity assays were performed by serial dilution of *E. coli* XL1-Blue strains onto LB-agar containing 200 μg/mL trimethoprim, 15 μg/mL gentamicin, and 0.1% (w/v) L-rhamnose. To test immunity protein function, bacteria were spotted onto LB-agar containing 200 μg/mL trimethoprim, 15 μg/mL gentamicin, 0.1% L-rhamnose, and 0.5 mM IPTG. *B. fragilis* strains used in toxicity assays were *B. fragilis* NCTC 9343 Δ*tdk* ΔBF9343_RS08070, *B. fragilis* NCTC 9343 Δ*tdk* ΔBF9343_RS08095, and *B. fragilis* NCTC 9343 Δ*tdk* ΔBF9343_RS08070 ΔBF9343_RS08095 (Supplemental Table 6). Strains were grown anaerobically on supplemented blood heart infusion agar plates containing gentamycin (60 μg/mL) and erythromycin (12.5 μg/mL) for 24 hours before cells are scraped and resuspended in PBS [42]. The OD_600_ was taken for each *B. fragilis* strain and adjusted to OD 0.1 prior to inoculation in BHIS broth containing either the gentamycin-only control (60 μg/mL) or containing anhydrotetracycline (aTC) 25 ng/mL. BHIS liquid media was deoxygenated for 2 hours prior to inoculation and cells were grown subsequently grown under anaerobic conditions. 6 hours after inoculation, bacterial cultures were pelleted, resuspended in 1 mL 1X phosphate buffered saline (PBS), and serially diluted by spotting on BHIS plates containing gentamycin (60 μg/mL). All assays were performed at least three independent times.

### Protein Expression and purification

For the expression and purification of effector-immunity complexes, 50mL of *E. coli* BL21 cells harboring the desired expression plasmid were grown overnight in LB to stationary phase. The next day, cultures were back-diluted 1:80 into 4L of LB media and grown to an approximate OD_600_ of 0.6 before protein expression was induced by the addition of 1 mM IPTG for three hours. Cells were then collected by centrifugation at 6000*g* for 15 minutes and frozen at −20C. For Ni-NTA purification, cells were thawed and resuspended in lysis buffer containing 50 mM Tris-HCl at pH 8.0, 150 mM NaCl, 10 mM imidazole, and subsequently lysed by sonication (6 x 30s pulses at 30% amplitude). Cellular debris was then cleared by centrifugation and bound proteins were eluted with buffer containing 150 mM NaCl, 50 mM Tris-HCl at pH 8.0, 400 mM imidazole. Protein samples were further purified by size exclusion chromatography using a HiLoad 16/600 Superdex 200 column on an ÄKTA FPLC system (GE Healthcare).

For individual immunity proteins, expression and purification were performed as described above except that following induction, cultures were incubated overnight at 18 °C to enhance protein expression. Reducing agents were included throughout purification to prevent intermolecular disulfide bond formation: 1 mM of dithiothreitol (DTT) was added to the lysis and elution buffers, and 1 mM β-mercaptoethanol (BME) was included in the SEC buffer. DTT was used during Ni-NTA affinity purification due to the incompatibility of BME with the resin.

For the purification of E2 effector, a slightly longer version of E2_tox_ was used (hereafter referred to as E2_CTD_) because it gave higher protein yields than E2_tox_. The E2_CTD_-I2 complex was then purified using a modified protocol to denature the immunity protein prior to Ni-NTA elution. Following binding of His-tagged E2_CTD_ in complex with untagged I2 to the column, the resin was washed with lysis buffer containing 6 M guanidinium chloride to remove I2. The column was subsequently washed three times with lysis buffer lacking denaturant to remove residual guanidinium chloride, and bound E2_CTD_ were eluted under native conditions. All purification steps were performed at pH 7.0 to preserve E2_CTD_ stability.

For isothermal titration calorimetry (ITC) experiments, E2_CTD_ and immunity proteins were further purified by size-exclusion chromatography (SEC) in the SEC buffer at pH 7.0 to ensure identical buffer conditions and eliminate heat artifacts arising from buffer mismatches.

### Size exclusion chromatography with multi-angle laser light scattering (SEC-MALS)

Size exclusion chromatography with multi-angle laser light scattering was performed on the E2_CTD_-I2 complex at a concentration of 2 mg/ml. The protein was further purified using a Superdex 200 column Increase 10/300 column (GE Healthcare). MALS data were collected using a MiniDAWN detector and Optilab system (Wyatt Technologies) and analyzed with the Astra software package (https://www.wyatt.com/products/software/astra.html).

### Isothermal Titration Calorimetry (ITC)

ITC measurements were performed with a MicroCal PEAQ-ITC microcalorimeter (Malvern). Titrations were carried out with 125 μM of the purified immunity protein in the syringe and 20 μM of E2_CTD_ in the cell. The titration experiment consisted of one 0.4-μl injection followed by 24 1.5-μl injections with 150-s intervals between each injection. The ITC data were analyzed using the Origin software (version 5.0, MicroCal, Inc.) and fit using a single-site binding model.

### Sequence Alignments and 3D-structure Prediction

All sequence alignments containing predicted secondary structure were performed using ESPript 3.0 [43]. For the prediction of three-dimensional protein structures and complexes, we used AlphaFold3 [44]. Figures were prepared with ChimeraX (1.8) [45].

### Western blot analysis

Western blots of protein samples were performed using an SDS-PAGE gel and buffer system and a standard western blotting protocol. After SDS-PAGE separation, proteins were wet-transferred to 0.45 μm nitrocellulose membranes (100 V for 30 mins, 4°C). The samples were then analyzed with Western blot using the protein-specific rabbit primary antibodies α-VSVG (1:5000, 1hr) and a goat α-rabbit secondary antibody (Sigma, 1:5000, 45 minutes). Western blots were imaged using a ChemiDoc System (Bio-Rad).

#### *E. coli* growth assays

Individual bacterial strains were grown overnight at 37°C in LB on a shaking incubator. The following day, 1 μL of the overnight culture was inoculated into 200 μL of fresh LB broth in a 96-well plate and incubated at 37°C with shaking. OD_600_ readings were taken every 15 minutes using an Epoch 2 plate reader (Biotek) until the OD_600_ of the strains were close to 0.3 at which point the expression of toxin and immunity proteins was induced via the addition of rhamnose and IPTG, respectively. Afterwards, the strains were allowed to grow under the same growth conditions for 15 more hours. For every given strain and condition, the experiment was independently repeated three times and the average growth is shown in the graph. The error bars represent the standard deviation of three biological replicates.

### Construction of inducible GA1 E2 effector plasmid and orphan immunity gene deletion mutant plasmid

All oligonucleotides used in this study are listed in Supplementary Table 6. The plasmid pNBU2_erm-TetR-P1T_DP-GH023_GA1_E2_CTD was created by amplifying the C-terminal domain of the effector GA1_E2 (HMPREF0999_RS10520) from *Parabacteroides* sp. D25 gDNA template with Phusion high-fidelity DNA polymerase (NEB) and cloned via Gibson Assembly into pNBU2_erm-TetR-P1T_DP-GH023 digested with BamHI-HF and SalI-HF restriction enzymes (NEB) using NEBuilder Hifi DNA assembly master mix (NEB). The plasmid pExchange-*tdk*_ΔBF9343_RS08070 was created by amplifying upstream and downstream homology regions flanking the orphan immunity gene (BF9343_RS08070) homologous to the cognate GA1_I2 from *B. fragilis* NCTC 9343 genomic DNA and cloned via Gibson Assembly into pExchange-*tdk* digested with BamHI-HF and Sal1-HF restriction enzymes. Plasmid sequences were confirmed via Sanger sequencing (Molecular Biology Shared Resources). The plasmid pExchange-*tdk*_ΔBF9343_RS08095 was created by amplifying upstream and downstream homology region flanking the orphan immunity gene (BF9343_RS08095) e2i2 pair from *B.fragilis* NCTC 9343.

#### *B. fragilis* genetic manipulation

Integration of pNBU2_erm-TetR-P1T_DP-GH023_GA1_E2_CTD plasmid into *att* sites was performed through mating between *E. coli* S17-1 λ *pir* and *B. fragilis* strains listed in Supplementary Table 6. For mating, a volume of 50 mL of *E. coli* was incubated in LB to late exponential phase (OD_600_ = 0.5) on an orbital shaker and a volume of 10 mL of BHIS media inoculated with *B. fragilis* was incubated to exponential phase under anaerobic conditions. Each culture was pelleted and washed prior to culture mixing before plating at high density on non-selective agar media and incubated for 16 hours at 37°C in an aerobic incubator. The cells were resuspended in 1 mL of BHIS and plated on BHIS plates containing gentamycin (60 μg/mL) and erythromycin (12.5 μg/mL). The insertions were verified by PCR (Supplemental Table 6) and att2 insertions were used in all cases.

For the generation of in-frame chromosomal deletions using pExchange-*tdk* plasmid and its derivatives, matings were performed as previously described above by between *E. coli* S17-1 λ *pir* and *B. fragilis* strains listed in Supplemental Table 6 After mating, merodiploid integrants were selected on BHIS plates containing gentamycin (60 μg/mL) and erythromycin (12.5 μg/mL), then were streaked on BHIS non-selective media, resuspended, and plated on BHIS plates containing floxuridine (200 μg/mL), and candidate deletion mutants were verified by PCR [46].

### Detection of genes in Bacteroidales genomes

We obtained a list of genera in the Bacteroidales order using the NCBI taxonomy browser, excluding candidatus, unclassified, and environmental genera. We downloaded the proteomes from species belonging to these genera from RefSeq on Feb 12, 2024 and created a combined blastp database. Using blastx we searched for homologs of effector, immunity and orphan immunity genes in this database that matched with a minimum of 90% protein identity along 90% of the length of the query gene. For GA1 effector genes we only searched with the final 399 bases of the C-terminal domain (CTD) were used due to its homology with other effector genes in the Rhs regions [10, 12]. For detection of orphan I2 genes in rAID systems from Bacteroidales reference genomes, we first identified rAID systems previously published methods [16]. Briefly, we identified rAID tyrosine recombinase homologs via high e-value scores from blast analysis using as our query the reference sequence from *B. fragilis* NCTC 9343 (BF9343_RS08045). We then assessed if homologs were found upstream of two or more co-directionally oriented genes with less than 41% GC content over their length, as previously defined. Hits meeting these criteria were defined as rAID systems and assessed for the presence of orphan I2 genes via blast.

### Metagenomic analysis

The species abundance in metagenomic samples was quantified using MetaPhlAn3 (version 3.0.14, parameters: read_min_len 30, database mpa_v30_CHOCOPhlAn_201901) [47]. Gene abundance was determined by mapping reads with Bowtie2 (version 2.3.4.1, parameters: -a -N 1), followed by normalizing read counts against gene length and library depth. The abundance of *B. vulgatus* single nucleotide polymorphisms (SNPs) was assessed from normalized counts of the four nucleotides at each site. Specifically, metagenomic reads were aligned to *B. vulgatus* MetaPhlAn3 marker genes using Bowtie2 (version 2.3.4.1, parameters: -a -N 1) [48].

These alignments were converted to pileup format using Samtools’ mpileup (version 1.17, parameters: --excl-flags UNMAP,QCFAIL,DUP -A -q0 -C0 -B) [49]. We then extracted nucleotide counts from the pileup, excluding the first and last 10 bases of each gene due to typically poor alignment quality, to calculate relative SNP abundances.

Abundance quantification of *P. vulgatus* SNPs was performed on both assembled genomes and metagenome alignments to *P. vulgatus* genes. For genomes, MetaPhlAn3 marker gene orthologs were identified using Prokka and BLAST, followed by gene alignment and variant identification [50]. SNPs were designated as strain markers if present in ≥90% of the strain’s genomes. In metagenomes, shotgun reads were aligned to target genes, searching for variants with a minimum coverage of 5, a minor allele count of at least 2, and presence in ≥10% of the samples. When examining the metagenome abundance of variants on the level of the individual gene, we found a single gene (A0A3E4WAR6) with an excess of SNPs whose abundance pattern was opposite other marker genes (variants found in the orphan encoding strain had a low abundance, whereas variants found in the 26 other marker genes had a high abundance). Because this gene was an outlier compared to other genes, it would be statistically irresponsible to include it when we averaged individual maker genes into an overall strain-level abundance, and therefore we excluded it from our calculations. It is like that there are closely related genes that are erroneously aligning in the metagenome.

### Phylogenetics

We developed a phylogeny for *P. vulgatus* strains from individual S01-am [25] using a methodology akin to our previously published work [42]. Firstly, Prokka (version 1.14) was utilized for gene identification, and blastn (version 2.12) to locate orthologs of each MetaPhlAn3 marker gene specific to *P. vulgatus*. Acceptance criteria for genes included a minimum of 90% identity and 90% coverage of the gene length. Additionally, genomes needed to contain at least 80% of these marker genes. Each marker gene was aligned individually using MAFFT (version 7.52) [51], followed by concatenation of these individual alignments. RAxML (version 8.2) [52] was then employed to construct a phylogenetic tree from the combined alignment. Visualization of this tree was facilitated using ETE3 [53], with an emphasis on indicating the presence of specific genes (GA1 E2, the canonical I2, two orphan variants of I2, orf3, and orf5) in each genome. The presence of these genes was determined using blastp against known references. However, for GA1 E2, only the final 399 bases of the C-terminal domain (CTD) were used due to its homology with other effector genes in the Rhs regions [10, 12].

### Positive selection analysis

We assessed selection in the metagenome-identified SNPs of orphan immunity orf3 and orf5 using two methods. Initially, we employed a sliding window approach to calculate the *dN/dS* ratio (non-synonymous to synonymous SNP ratio) along both genes. This was done using 200 base pair sliding windows, in increments of 25 base pairs. In each window, the *dn/ds* ratio was calculated, incorporating a background count of 1 to prevent division by zero, analogous to a Beta(1,1) prior for the estimator. Regions were considered under positive selection if their *dn/ds* ratio exceeded 3. Additionally, we identified a second type of selection based on a statistically significant temporal change in SNP abundance. This significance was determined using the p-value obtained from linear regression analysis.

### Strain calling from metagenomic data

We identified strains of *P. vulgatus* in metagenomic data using StrainFacts [54]. We obtained a matrix of nucleotide counts along *P. vulgatus* metaphlan markers described above. To ensure that StrainFacts ran in an acceptable amount of time, we focused on bases that had at least a coverage of 5 and at least 2 counts of the minor allele in at least 10% of samples. We ran StrainFacts with a gamma hyperparameter of 10-20 to force the algorithm towards discrete, non-probabilistic genotypes. We chose the number of strains with a number that gave us metagenomic strains that correspond to two strains we found at the genome level, one encoding the cognate genes and one encoding the orphan genes. Thus, we took the minimum number of strains required to achieve a reasonable correlation (>0.3) to both the cognate and orphan genes. We find that this occurs with 4 strains (correlation of 0.42 to both). We reconstructed the sequences for each strain by inserting variant bases that could be assigned with some confidence to the reference or alternate allele (probability < 0.25 or > 0.75), otherwise inserting an unknown base. Reconstructed sequences were treated the same as the sequences as orthologs from genomes and a new tree was constructed using the methods above. We identified the number of strains by the principle of non-redundant strains: strains that have highly correlated abundance over time are redundant and add nothing new to our understanding of the microbiome strains. In detail, for a fixed number of strains we consider the highest strain-strain abundance correlation and we select the number of strains that minimizes that value.

---

**Supplemental Figure 1. Example rAID systems.** Selected rAID systems harboring orphan I2 genes from reference genomes of Bacteroidales.

**Supplemental Figure 2. Alignment of GA1_I2 amino acid sequences. A)** Amino acid sequence alignment of cognate GA1_I2 and orphan I2 genes from the rAID system of *B. fragilis* NCTC 9343. **B)** Amino acid sequence alignment of cognate GA1_I2 sequences from *B. uniformis* and *P. vulgatus* (BU and PV), rAID orphan immunity from S01-am (orf3 and orf5), and the single domain SUKH protein from *Methylomonas sp.* LW13 (Meth).

**Supplemental Figure 3. Western blot assaying for the expression of I2^5x^**. Western blot demonstrating equivalent levels of protein in lysates from *E. coli* cells that express WT and mutated (I2^5x^) proteins using anti-VSVG.

**Supplemental Figure 4. Abundance plots of marker genes detected in longitudinal samples from S01-am.**

**Supplemental Table 1. GA1 T6SS E2 and I2 homologs detected in reference genomes.**

**Supplemental Table 2. GA1 T6SS I2 homologs identified in the Integrated Gene Catalog.**

**Supplemental Table 3. GA1 T6SS I2 homologs identified in rAID systems from reference genomes of Bacteroidales.**

**Supplemental Table 4.** GA1 T6SS E2 and I2 homologs identified in longitudinal isolate genomes from Poyet *et al*.

**Supplemental Table 5. Amino acid altering SNPs detected in orphan I2 genes in S01-am metagenomes.**

**Supplemental Table 6. Strains, plasmids, and primers used in this study.**

