## Supplementary figures and images for "Rapidly evolving orphan immunity genes protect human gut bacteria from intoxication by the type VI secretion system"

### Supplemental Figures 1-4

#### Supplemental Figure 1

A

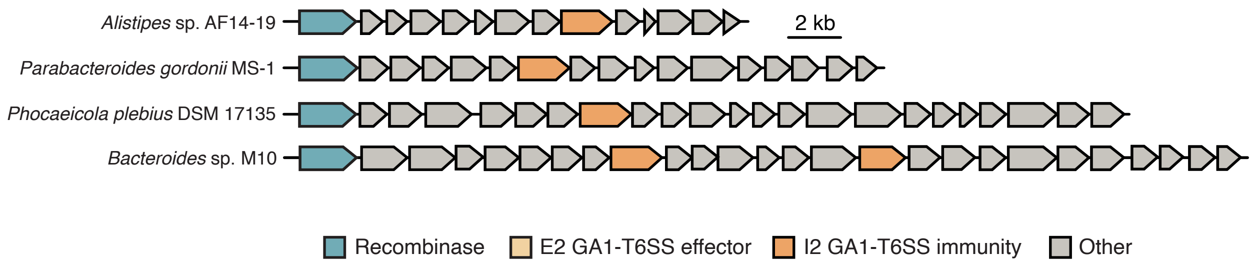

#### Supplemental Figure 2

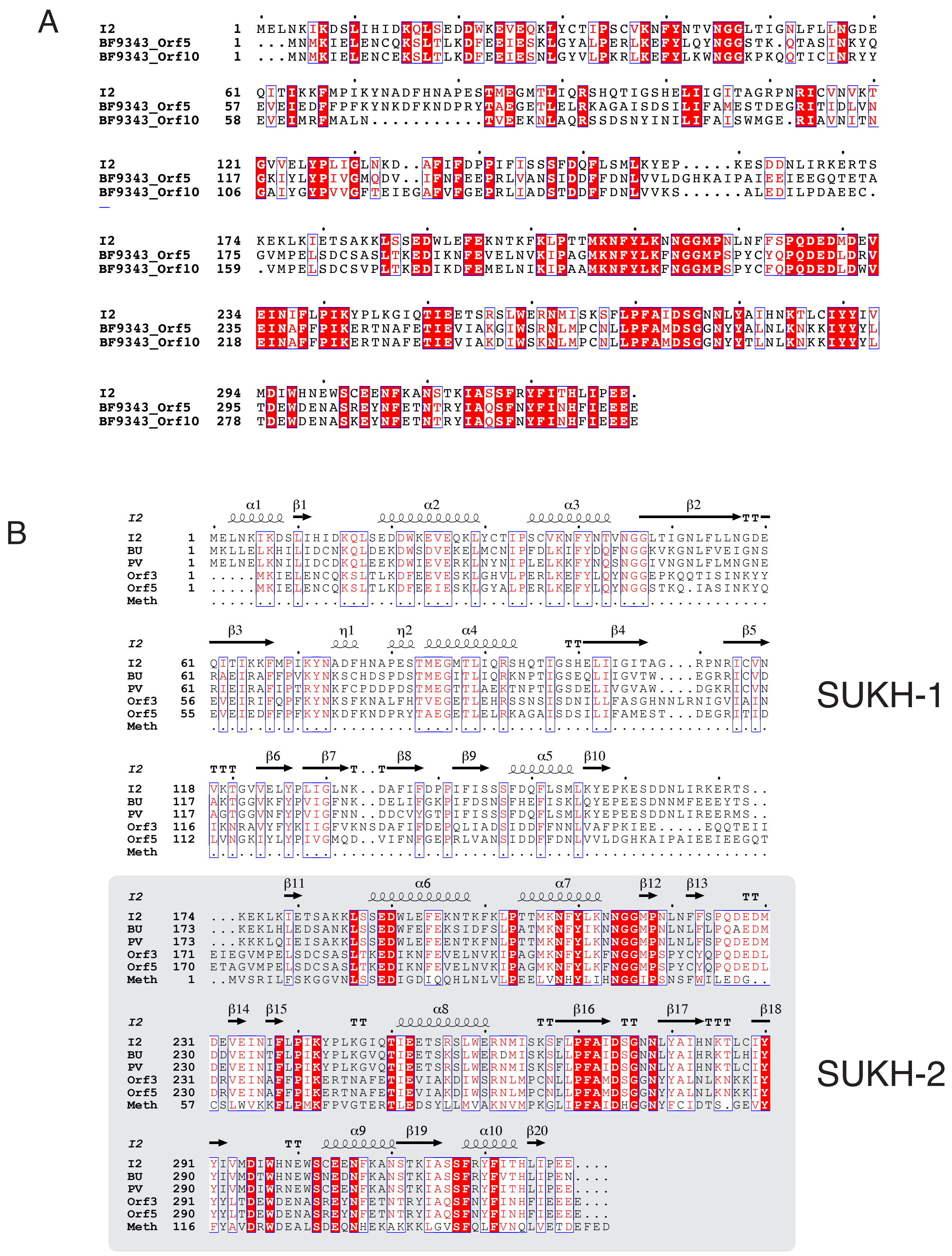

#### Supplemental Figure 3

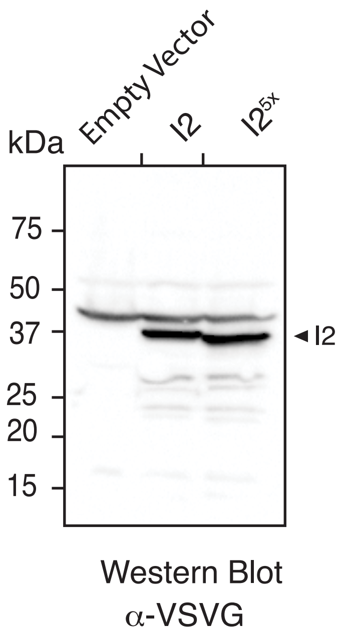

#### Supplemental Figure 4

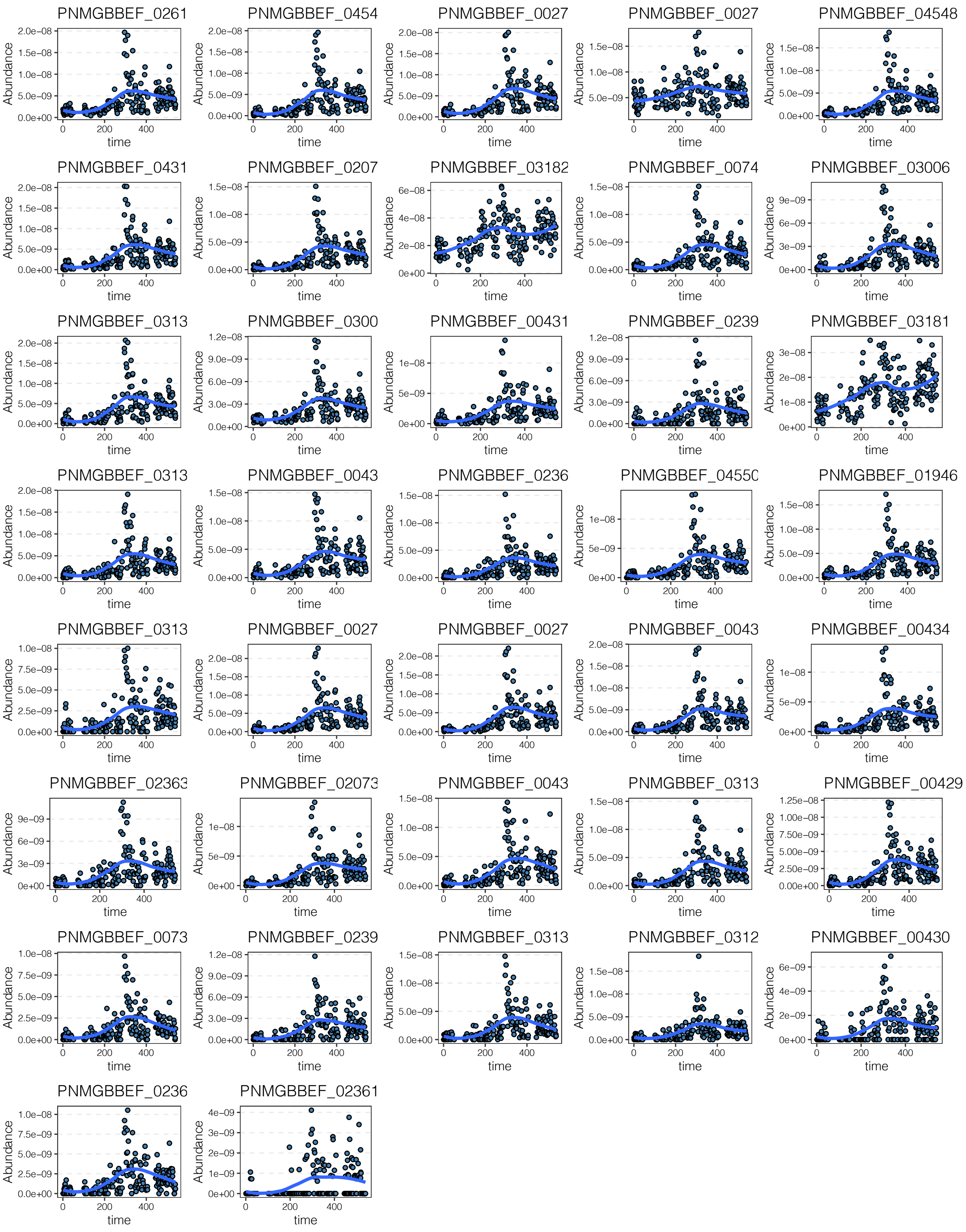
